# On the mechanistic roots of an ecological law: parasite aggregation

**DOI:** 10.1101/680041

**Authors:** Jomar F. Rabajante, Elizabeth L. Anzia, Chaitanya S. Gokhale

## Abstract

Parasite aggregation, a recurring pattern in macroparasite infections, is considered one of the “laws” of parasite ecology. Few hosts have a large number of parasites while most hosts have a low number of parasites. Phenomenological models of host-parasite systems thus use the negative-binomial distribution. However, to infer the mechanisms of aggregation, a mechanistic model that does not make any a priori assumptions is essential. Here we formulate a mechanistic model of parasite aggregation in hosts without assuming a negative-binomial distribution. Our results show that a simple model of parasite accumulation still results in an aggregated pattern, as shown by the derived mean and variance of the parasite distribution. By incorporating the derived statistics in host-parasite interactions, we can predict how aggregation affects the population dynamics of the hosts and parasites through time. Thus, our results can directly be applied to observed data as well as can inform the designing of statistical sampling procedures. Overall, we have shown how a plausible mechanistic process can result in the often observed phenomenon of parasite aggregation occurring in numerous ecological scenarios, thus providing a basis for a “law” of ecology.

## Introduction

Parasites are ubiquitous [1]. Nevertheless, for parasitologists, sampling can be a hard problem [2, 3]. This is due to the distribution of parasites among hosts [4]. While most hosts harbour very few parasites, only few individuals are hosts to a large number of parasites [5]. Normal distribution is not appropriate to represent such parasite distribution in a host community. This phenomenon, termed as parasite aggregation, is perhaps one of the few “laws” in biology because it is a recurring pattern in nature, and finding exceptions to this pattern is rare [6, 7, 8]. The pattern is specifically observed in macroparasite infections, which include diseases brought about by helminths and arthropods [7, 8]. Knowing the macroparasite distribution in host populations is essential for addressing challenges in investigating disease transmission. Examples are human onchocerciasis and schistosomiasis [9, 10, 11]. Aggregation can also affect co-infection by various parasites, parasite-driven evolutionary pressures, and stability of host-parasite communities [12, 13, 14, 15]. Thus, the study of parasite aggregation addresses a fundamental issue in ecology.

The distribution of hosts with different numbers of parasites can be well captured by the negative-binomial [5, 16, 17]. Given this fact, several theoretical models about host-parasite dynamics implicitly assume the negative-binomial distribution, mainly based on a phenomenological (statistical) modelling framework [18, 6, 8, 19]. The phenomenological principle is based on the observation that - (i) a distribution is Poisson if showing random behaviour with equal mean and variance, (ii) binomial if under-dispersed with higher mean than variance, or (iii) negative-binomial if overdispersed with higher variance than the mean [12, 20]. Overdispersion is observed in host-parasite interaction, such as between cattle (*Bos taurus*) and warble fly (*Hypoderma bovis*), and between Nile tilapia (*Oreochromis niloticus*) and copepod *Ergasilus philippinensis* [12, 21]. The negative-binomial distribution is hypothesized to arise due to the heterogeneity in the characteristics of the host-parasite interaction, and environmental variation. Heterogeneous exposures, infection rates and susceptibility of host individuals can produce aggregated distributions of parasites [12, 22, 23, 24]. However, is heterogeneity in the characteristics of the hosts and parasites a sufficient condition for aggregation [25]? Using a mechanistic model, we show that a regular but non-extreme parasite accumulation can lead to aggregation, and neither to Uniform nor Poisson distribution. We show that even in regular stochastic host-parasite systems (i.e., with homogeneous conditional probabilities of macroparasite accumulation), aggregation is widely possible.

Traditionally, in arriving at the desired model for host-macroparasite interaction, phenomenological modelling is assumed [18, 26, 5, 27, 28]. Phenomenological models are based on data gathered, but typically, the mechanisms underlying the phenomenon are hidden. There is a need for descriptive and predictive models that assume the underlying mechanistic processes of parasite dynamics, useful in formulating disease control programs [2]. The goal of this study is to mechanistically illustrate the interaction between hosts and macroparasites without assuming a negative-binomial distribution. Our approach considers parasite accumulation without direct reproduction in host individuals, which is a consequence of the complex life history of macroparasites [29, 30, 31]. For example, the Nile tilapia fish are infected by the acanthocephalans parasites via the ingestion of infected zooplankton. The parasites grow, mate and lay eggs inside the fish, and then eggs are expelled in the lake through faecal excretions. The parasites in the excretions are passively consumed by the intermediate hosts (zooplanktons), and the cycle of infection through foraging continues [32, 33].

### 1.1 The gap in existing models

Recently, various attempts have been made to model the accumulation and aggregation of parasites mechanistically. The stratified worm burden, which is based on a chain of infected compartments as in Susceptible-Infected (S-I) modeling framework, was used to model schistosomiasis infections [34, 35, 36]. This model can be well simulated numerically for specific cases, but possibly difficult to implement analytic studies to find general mathematical conclusions [34]. Although there are several studies that have considered addressing this issue (e.g., simplifying infinite dimensional differential equations) [37], our model proposes an alternative and simple approach.

The Poisson-Gamma Mixture Model has been proposed to model aggregation based on parasite accumulation [38]. The negative-binomial distribution arises naturally from a Poisson-Gamma process; however, is it possible to derive a mechanistic model of aggregation without initially assuming a distribution related to the negative-binomial? The answer to this question is “yes”, and there are existing studies that consider bottom-up mechanistic models [39, 37, 40]; however, these models are very few compared to the phenomenological approaches available. Our model is proposed as an alternative approach, but with unique features, that can be added to the list of available models useful for studying diverse types of macroparasites. One such feature of our model is that we can predict over a wide-range of parameter space when underdispersed and random parasite distributions (not just the overdispersed distribution) will emerge. Our model also predicts the emergence of the Geometric distribution, confirming previous predictions [41], which can arise even in the absence of heterogeneity.

In another study, a mechanistic model has been proposed based on the random variation in the exposure of hosts to parasites and in the infection success of parasites [6]. This model anchors its derivation and conclusion on the common assumption that heterogeneity in individual hosts and/or parasites leads to parasite aggregation. In our proposed model, we can have further insights on the resulting distribution of parasites given homogeneous or heterogeneous biological, ecological and environmental factors affecting the host-parasite interaction.

Moreover, in most models of macroparasite infections, the rate of disease acquisition by susceptible individuals (known as the “force of infection” *β*) is assumed as a parameter. However, identifying the value of *β* can be difficult or infeasible even in the availability of data [29]. In our proposed model, this challenge can be addressed since our assumed infection parameter, denoted by *P*, can be directly estimated from parasite count data gathered from host samples in empirical studies.

Our model provides an alternative framework which is simple and tractable. Besides being useful in estimating required sampling size in the field, our work can be incorporated as part of classical modeling frameworks (e.g., host-parasite interaction with logistic growth) [42].

## Proposed model

The core concept of our model is captured in Fig. 1. The total host population has a density of *X*. With probability *P*_0_, the hosts are parasite-free, and the density of parasite-free hosts is then *X*_0_ = *P*_0_*X*. Similarly, the density of hosts which have at least one parasite is *X*_1_ = *P*_1_*X*. The total host population is therefore equivalent to *X* = *X*_0_ + *X*_1_ = *P*_0_*X* + *P*_1_*X*. Moreover, the class of individuals *X*_*i*_ which have at least *i* > 1 number of parasites is assumed as a subset of *X*_1_ (i.e., *X*_*n*+1_ ⊆ *X*_*n*_ ⊆ … ⊆ *X*_2_ ⊆ *X*_1_), and can be modeled as,

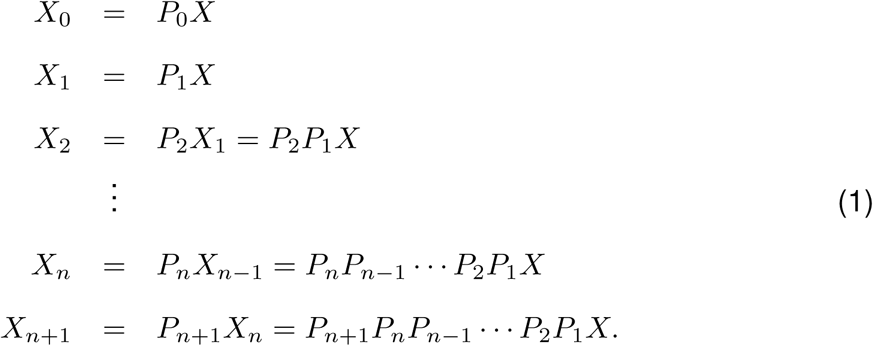

**Figure 1:**
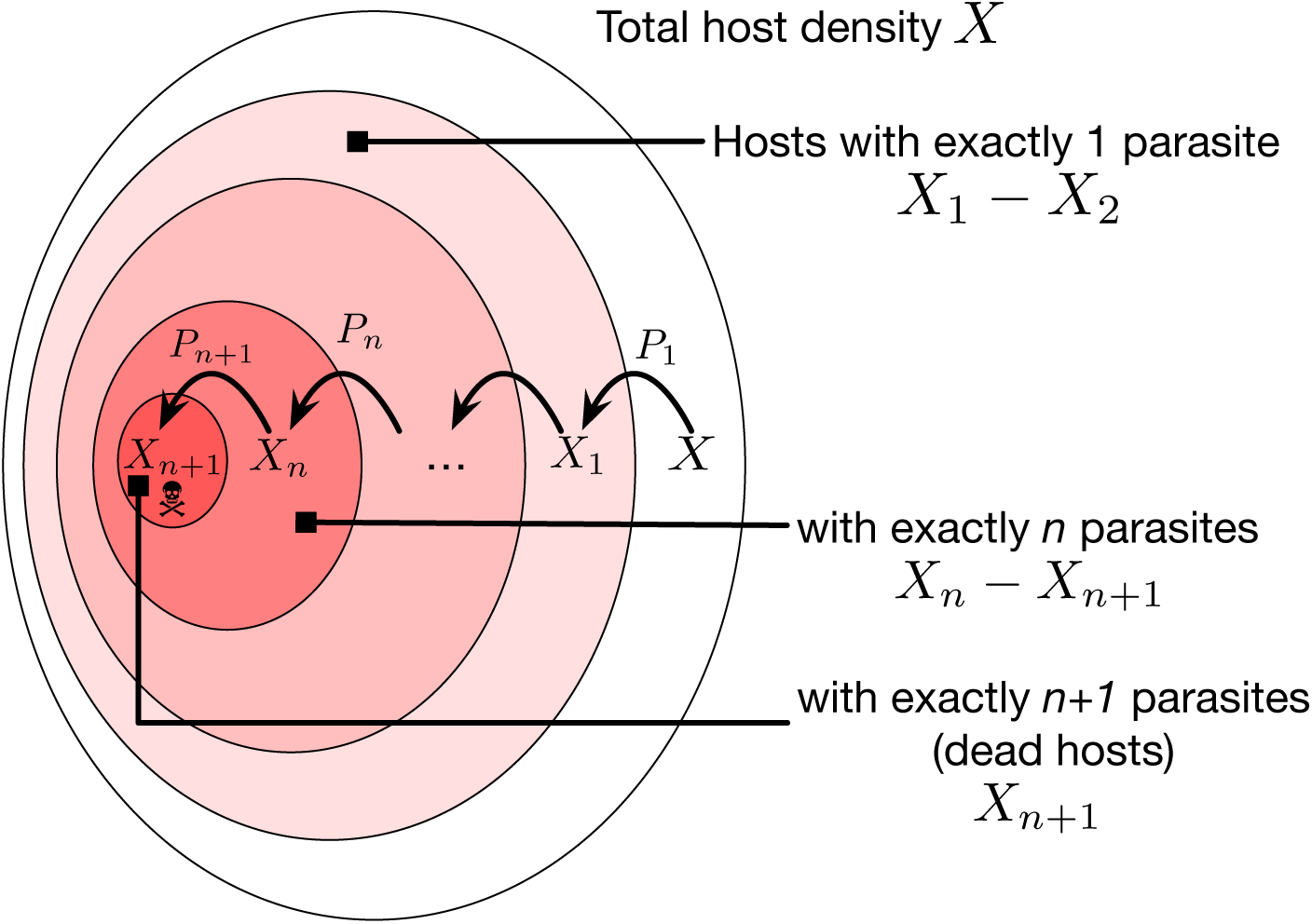
Hierarchical compartment model of macroparasite accumulation in hosts. The total host density is *X*. The density of parasite-free hosts is denoted by *X*_0_. The population density of hosts with *at least i* number of parasites is *X*_*i*_, *i* ≥ 1 (*X*_*i*+1_ ⊆ *X*_*i*_). A living host can harbour a maximum of *n* parasites. The parameter *P*_*i*_,*i* ≥ 1 can be interpreted as the net probability of parasite acquisition by a host in state *X*_*i*−1_. Refer to the Table in SI for the description of the state variables and parameters.

From the above set of equations, we can derive the distribution of living hosts (all except the *n* +1 compartment) with different parasite load concerning the total host population. Fig. 1 represents the classification of the states of the hosts depending on their parasite load. The compartment associated with *X*_*i*_, *i* ≥ 1 contains the hosts infected by at least *i* number of parasites. The parameter *P*_*i*+1_ characterizes the net probability that a host in *X*_*i*_ acquires an additional parasite (that is, probability of catching a parasite less the probability of removing a parasite from the host, e.g., through treatment or parasite death). Consequently, the difference *X*_*i*_ − *X*_*i*+1_,*i* ≥ 1 is the population density of hosts with *exactly i* number of parasites. The population density of living hosts (with maximum tolerable parasite load *n*) is then *X*_*n*_ − *X*_*n*+1_. Thus the death of hosts due to the parasite is captured by the transition into the *X*_*n*+1_ state given by 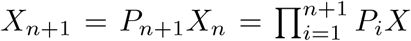, the number of dead hosts (last compartment in Eqs. (1)).

Our model, described as such, does not follow the classical input-output modelling framework (e.g., stratified worm burden model, which is an extension of the standard S-I epidemiological models). Such a classical modelling framework could be intractable when modelling aggregation [34]. The number of variables and equations in the stratified worm burden model can increase as the number of states (compartments) increases. Here, we model the states in the compartment diagram using proportions so we can efficiently analyze the distribution of parasites using probabilities. The dynamics of parasite infection can then be summarized using the properties of the derived distributions, such as the mean and variance. Also, in input-output models, the transfer from one state (e.g., susceptible) to another (e.g., the state with three parasites) needs to pass through intermediate diseased states (e.g., states with 1 and 2 parasites, respectively). In our model, a host can acquire more than one parasite, captured by adjusting the parameter *P*_*i*_. This approach is consistent with the experimental approaches and comparable to the data gathered. For example, the proportion 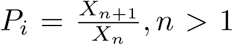 is directly computable from parasite load data from sampled hosts as compared to computing the force of infection *β* in S-I models [43].

The quantities relating to the host and parasite proportions, *X*_*i*_, *P*_*i*_ and *n* can be dynamic (e.g., may change over time). For example, we assume that *X*_*i*_ changes following a logistic growth with parasitism, and *P*_*i*_ is a function of parasite population *Y* (hence, also a function of time). This temporal connection allows us to relate our model to experimental data. The parameter values (e.g., *P*_*i*_) can be determined from the samples gathered at a specific time instant, and the pattern of the temporal evolution of the parameter values can be inferred from time-series data.

Depending on the exact transmission mode of the parasite, *P*_*i*_ can have different functional forms. An underlying assumption would be that *P*_*i*_ is a function of parasite encounter and transmission rates. For example, we could have 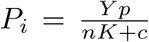 for all *i* ≠ 0. Here, the total ecological carrying capacity for the parasite population is assumed to be *nK* + *c*, where *K* is the host carrying capacity and *n* is the maximum number of parasites that a living host can harbour without dying. The parameter *c* is the quantitative representation of the environment where the parasites can survive outside the hosts. Thus we can interpret 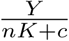 as the parasite encounter probability with *p* being the parasite transmission probability. We focus on cases where parasites highly depend on the hosts to survive, e.g., we assume a small value of *c* = 1. Homogeneity in parasite transmission is implemented by keeping *p* constant.

For the parasite population, the total parasite density in host population is 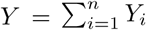 where *Y*_*i*_ = *i*(*X*_*i*_ − *X*_*i*−1_),*i* ≥ 1. *Y*_*i*_ is the population density of parasites associated with hosts having exactly *i* number of parasites. The distribution of parasites is according to the following:

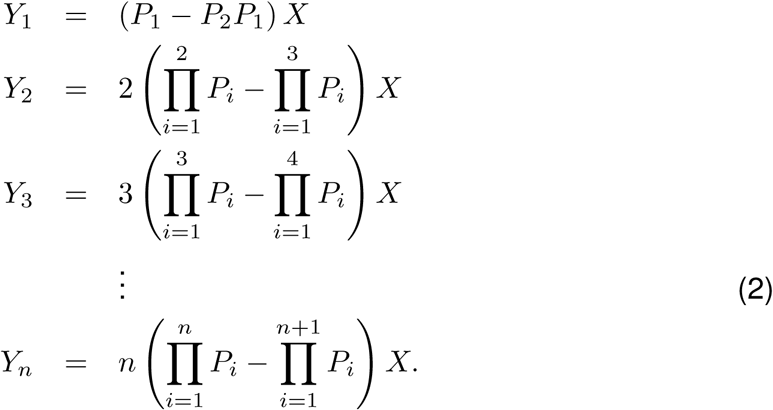

Based on the set of Eqs. (2), the parasite population density in host population can be written as 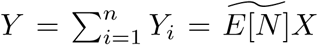 Here, *N* is the random variable representing the parasite load in a living host, and 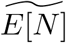 is the approximate mean of *N*.

### 2.1 Application to host-parasite population dynamics

The host basal growth rate is set to a constant *r*_*H*_, assuming that macroparasites do not affect reproduction of hosts. The carrying capacity of the host population is assumed to be equal to *K*. The death rate of the hosts is assumed to be 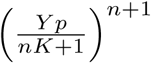, which is derived from the equation representing *X*_*n*+1_. Moreover, the per capita parasite reproduction rate is assumed to be *r*_*p*_. Together with the carrying capacity for the parasites (*nK* + 1), the population dynamics between the hosts and parasites can be modelled assuming logistic growth as follows (refer to Table SI.1 in SI for the description of the state variables and parameters):

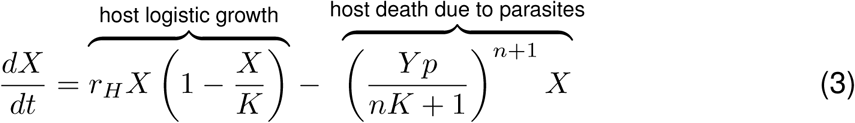

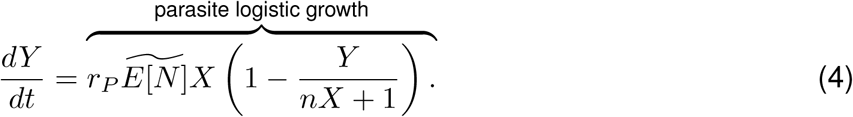

## Results

The distribution of the parasites in the living hosts is represented by 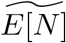. The general expression for the approximate mean of this distribution as discussed above is given by,

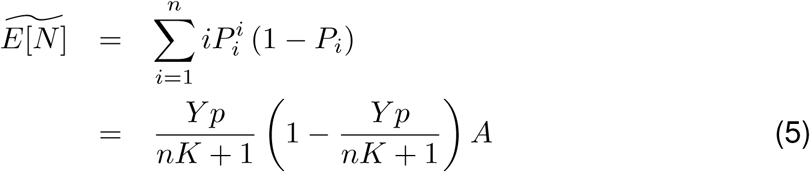

where *A* is derived from the derivative of a geometric series (see SI). If the parasite transmission 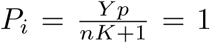 (for *i* ≠ 0), then the distribution of the parasites in the living hosts has 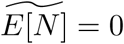 since all hosts that are harbouring at least *n* +1 number of parasites are dead. Now, supposing, 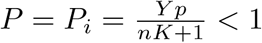, we have,

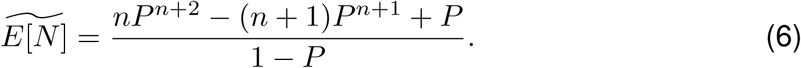

as the mean; and the variance, 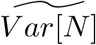, derived and presented in the SI.

A negative-binomial distribution describing the probability distribution of the number of successes before the *m*-th failure, where *ρ* is the probability of success can be written as *NB*(*m, ρ*). The mean and variance of *NB*(*m, ρ*) are 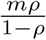 and 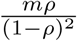,respectively. From Eq. (6), 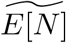 is equivalent to the mean of a negative-binomial distribution with *m* = *nP*^*n*+1^ − (*n* + 1)*P* ^*n*^ +1 and *ρ* = *P*. For the non-truncated negative-binomial distribution, we consider *n* → ∞ [5, 44]. In the next section (3.1), we discuss that as 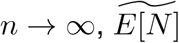 and 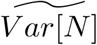 respectively converge to the mean and variance of a geometric distribution (*NB*(1, *ρ*)).

### 3.1 Constant host-parasite encounter probability

Suppose the parasite encounter probability 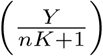 is fixed. This implies that what-ever the values of *Y* and 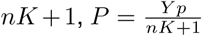 is always constant. We can also interpret *P* as the constant geometric mean of the parasite acquisition probabilities *P*_*i*_, *i* ≥ 1.

For a large *n* and 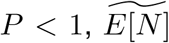 and 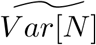 approximate the mean and variance of *N* ∈ {0, 1, 2, …, *n*}, respectively. *E*[*N*] can be interpreted as the average number of parasites in a host with variance *Var*[*N*]. From Eq. (6) as 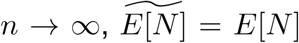 is equivalent to the mean of a negative-binomial distribution with *m* = 1 and *ρ* = *P*, which characterizes the mean of a geometric distribution. Let us denote this mean and variance as

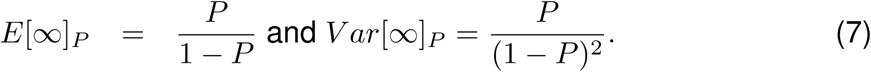

The variance-to-mean ratio is 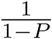, which increases as *P* increases (Table SI.2 in SI).

This result implies that even if the host can sustain a large number of parasites (*n* → ∞) for *P* < 1, we have finite mean and variance at the population level. Also, the variance-to-mean ratio is greater than 1 characterizing an overdispersed distribution of parasite load in the host population. For example, *E*[∞]_0.5_ implies that the average number of parasites in a host is 1 with variance *Var*[∞]_0.5_ = 2. *E*[∞]_0.9_ implies that the average number of parasites in a host is 9 with variance *Var*[∞]_0.9_ = 90.

As stated earlier, *E*[∞]_*P*_ is an approximation. The error due to the approximation can be estimated, shown in Fig. 2A. For a wide range of parameter values of the maximum tolerable parasite load of the host (*n*) and of the probability of parasite acquisition by a host (*P*), we see that the mean of the geometric distribution is a reasonable estimate for 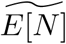. Fig. 2A further illustrates that as *n* approaches infinity, the error becomes smaller. While a small value for *n* and a higher *P* produce a higher error, this could also be an improbable scenario in nature. The condition where there is a high probability of acquiring more parasites but the host can only tolerate low parasite burden would result in morbid infection rates and potentially lead to the mortality and eventual extinction of the hosts. This possibility is illustrated in Fig. 3 where high parasite-driven host death drives the host population extinct 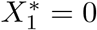.

**Figure 2:**
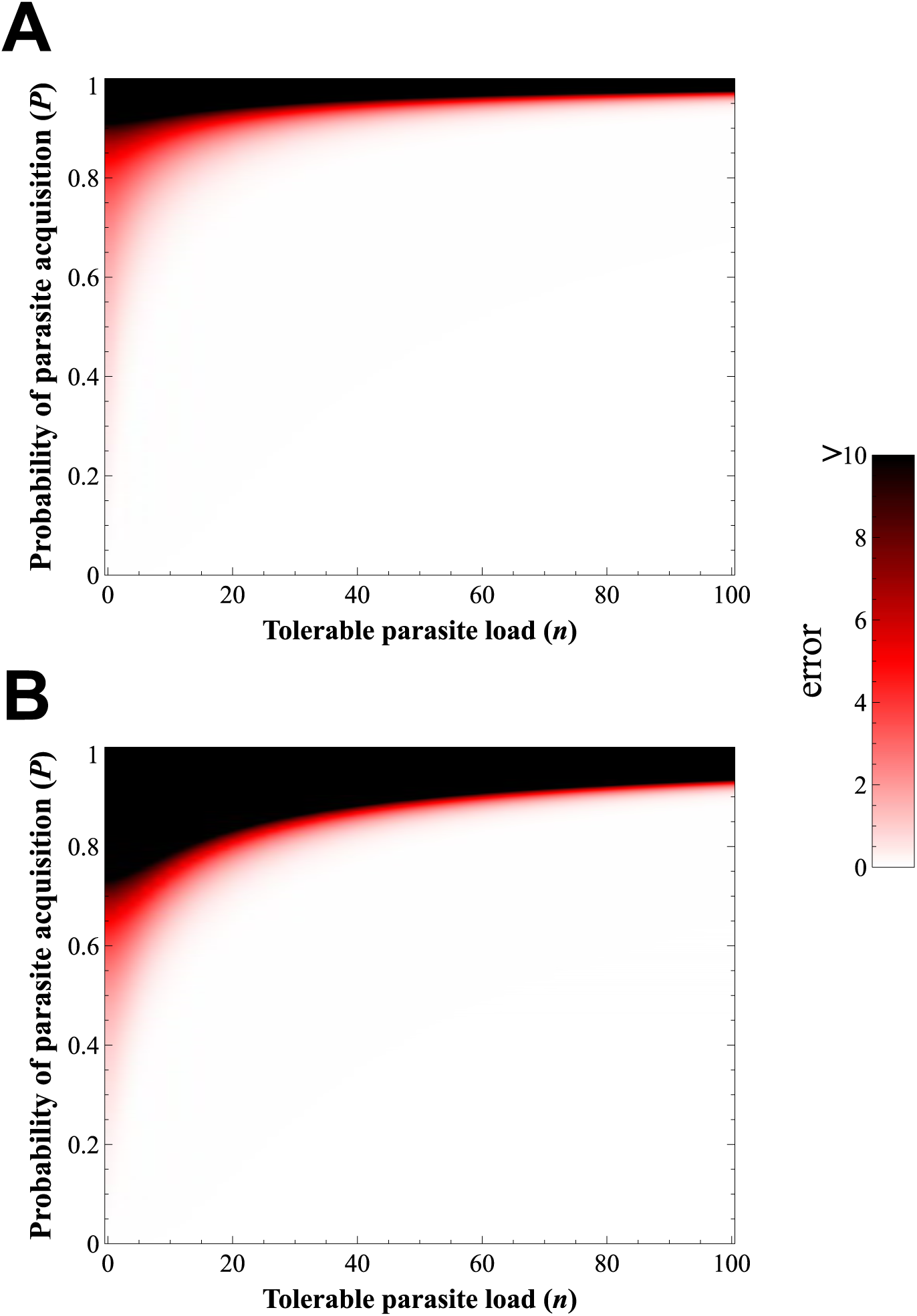
Approximation of 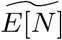 and 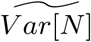 using the mean and variance of the geometric distribution, respectively. This shows how well the geometric distribution represents the parasite load distribution in hosts, especially when *n* is high and *P* is relatively low. (A) Absolute difference between 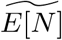 and 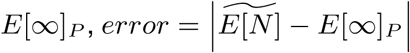. (B) Absolute difference between 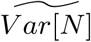 and 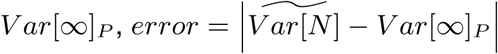.

**Figure 3:**
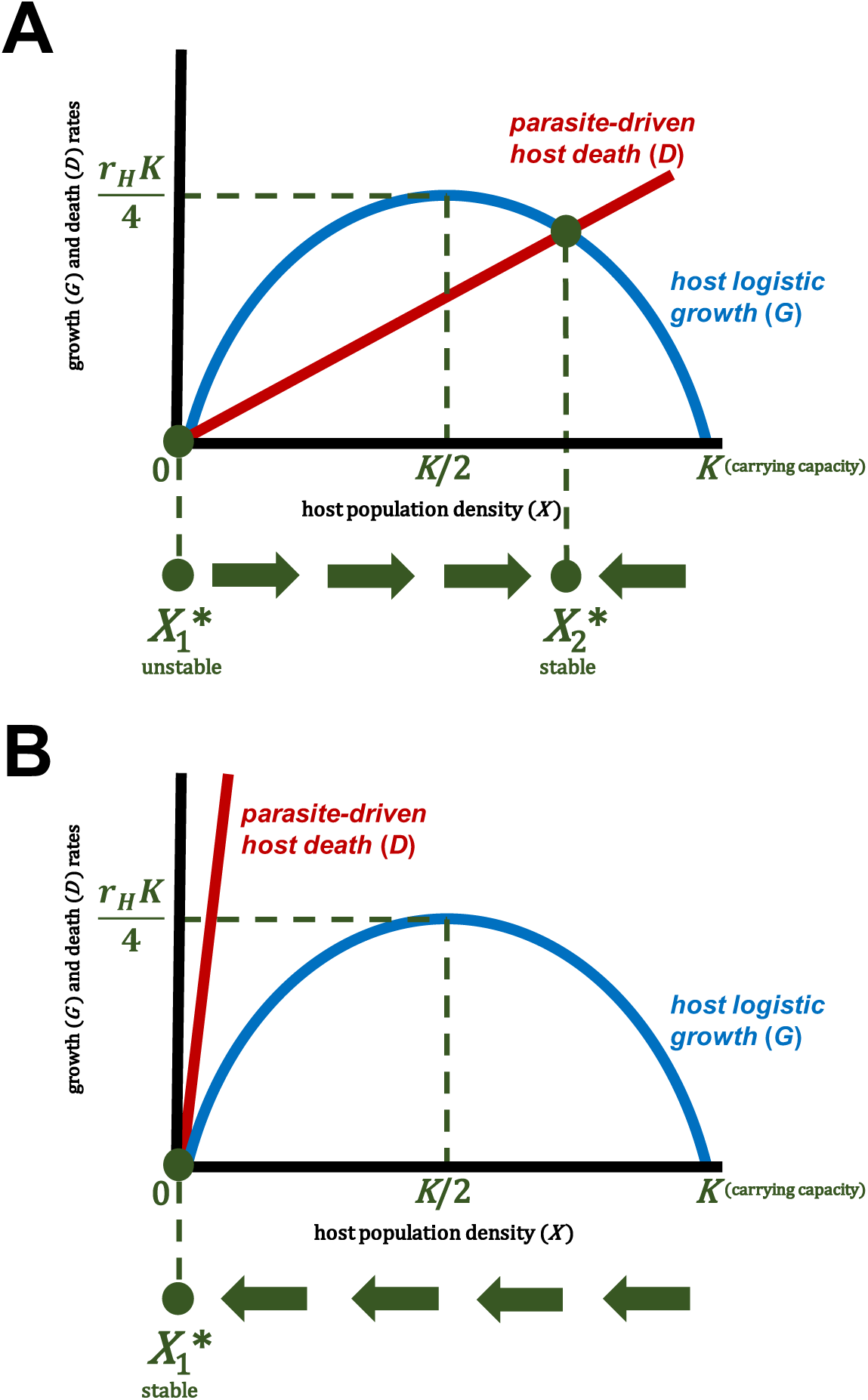
Illustration of host population dynamics. The population dynamics of hosts is based on Eq. (3). Suppose the host logistic growth function is 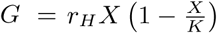 (blue curve) and parasite-driven death rate function is *D* = *P*^*n*+1^*X* (red line). The possible maximum growth rate of the hosts is *r*_*H*_*K/*4 where *r*_*h*_ is the host basal growth (reproduction) rate and *K* is the carrying capacity of the host population *X*. If *G* > *D* (blue curve is above the red line) then the population density of host increases. If *G*<*D* (blue curve is below the red line) then the population density of host decreases. An intersection of the growth curve (blue) and the death rate line (red) is an equilibrium point. (A) There are two equilib1r3ium points: the unstable 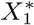 and the stable 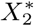. (B) There is one equilibrium point: the stable 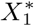 that represents parasitism-driven extinction of hosts. The red line with steep slope which possibly leads to the extinction of hosts and parasites can arise due to low value of *n* and/or large value of *P*.

The same is true with the approximation of 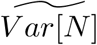 by *Var*[∞]_*P*_. Fig. 2 B shows that as *n* → ∞, the error becomes smaller. This result illustrates that the variance of the geometric distribution is a good estimate for 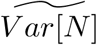 (Fig. 3).

In the supplementary information (Figs. SI.1 and SI.2), we present examples of parasite load distribution in the host population with differing values of *P*. If the value of *P* is intermediate, it results in aggregated distribution. However, the distribution becomes more negatively skewed as *P* increases which shows high host mortality due to harbouring high parasite load. Therefore, as *P* increases, the errors in approximating 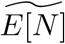 and 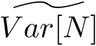 using the geometric distribution also increase.

#### 3.1.1 Population dynamics

We now analyze in detail the population dynamics between the hosts and parasites (Eqs. (3) and (4)). Since the probability of parasite acquisition by a host *P* is constant, we can decouple the host-parasite dynamics. Analyzing first the host population (Eq. (3)), we have a logistic growth function with death or harvesting term (*D* = *P*^*n*+1^*X*) [45]. In Fig. 3, the blue curve represents the logistic growth function *G*. The red line represents the death function *D*. The intersection of the two curves is an equilibrium.

There are two equilibrium points if *r*_*H*_ > *P*^*n*+1^, where one is unstable 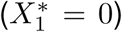 and the other is stable 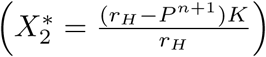 (Fig. 3A). If *r*_*H*_ ≤ *P*^*n*+1^, the zero equilibrium point is stable and the only steady-state of the dynamics (Fig. 3B), implying an eventual extinction of the host population. For the parasites to exploit the maximum growth rate of the hosts (*r*_*H*_ *K/*4) (Fig. 3), the parasite-driven host death rate should be *P*^*n*+1^ = *r*_*H*_ */*2.

Now, analyzing the parasite population (Eq. (4)), there are two possible equilibrium states: the parasite can go extinct, 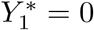 which happens if *X* = 0 or 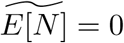; and a stable coexistence of hosts and parasites at 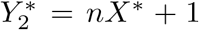 where *X*\* is the host equilibrium. The equilibrium state 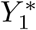 is unstable, and 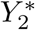 is stable. The condition 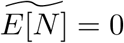 for 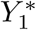 only happens if *P* = 0 (proof in SI).

Over time the total parasite population density in living host population converges to 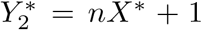. The expected number of parasites in the hosts then tends to *n* (excluding parasites in the environment represented by *c*, since they are outside the living hosts). One might think that this limiting case may not be the situation if parasite distribution in living hosts is aggregated. If aggregation affects the carrying capacity of the parasite population, the parameter *n* in the denominator *nX* +1 in Eq. 4 can be replaced by

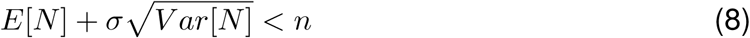

where *σ* represents the contribution of the variance to the average number of parasites in a host. Hence, if the distribution associated with parasite aggregation is considered, 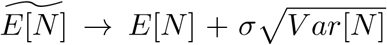 as time *t* → ∞. In the next section, we investigate the case when *P* is not constant through time.

### 3.2 Variable host-parasite encounter probability: Population dynamics

Until now, we assumed that the parasite-encounter probability *P* as a constant. However, 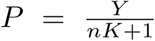 (Eqs. (3) and (4)) is a function of the dynamic variable *Y*, the parasite density. Given the dynamics of the parasite, we can have three possible equilibrium points. Two of the equilibrium points are trivial, one where no host and parasite exist (*X*\* = 0,*Y* * = *Y*_0_), and another at the carrying capacity of the hosts (*X*\* = *K, Y* * = *Y*_0_ = 0) with 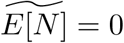 representing the disease-free state.

The third equilibrium point posits a coexistence of hosts and parasites and can be derived from

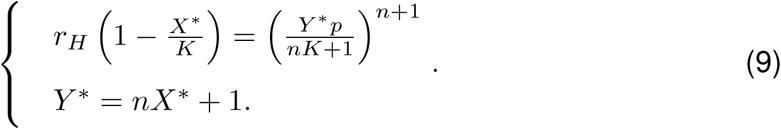

This leads to

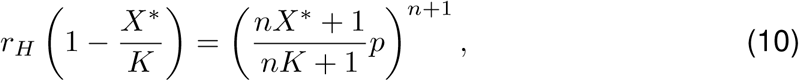

which we can analyze by investigating the intersection of the curves formed by the left and right hand sides of this equation. Suppose this intersection is *X*\* = *α* > 0 (Fig. 4). The equilibrium point is then (*X*\* = *α, Y* * = *nα* + 1) where *α* satisfies Eq. (10). This equilibrium point is stable (Fig. 4).

**Figure 4:**
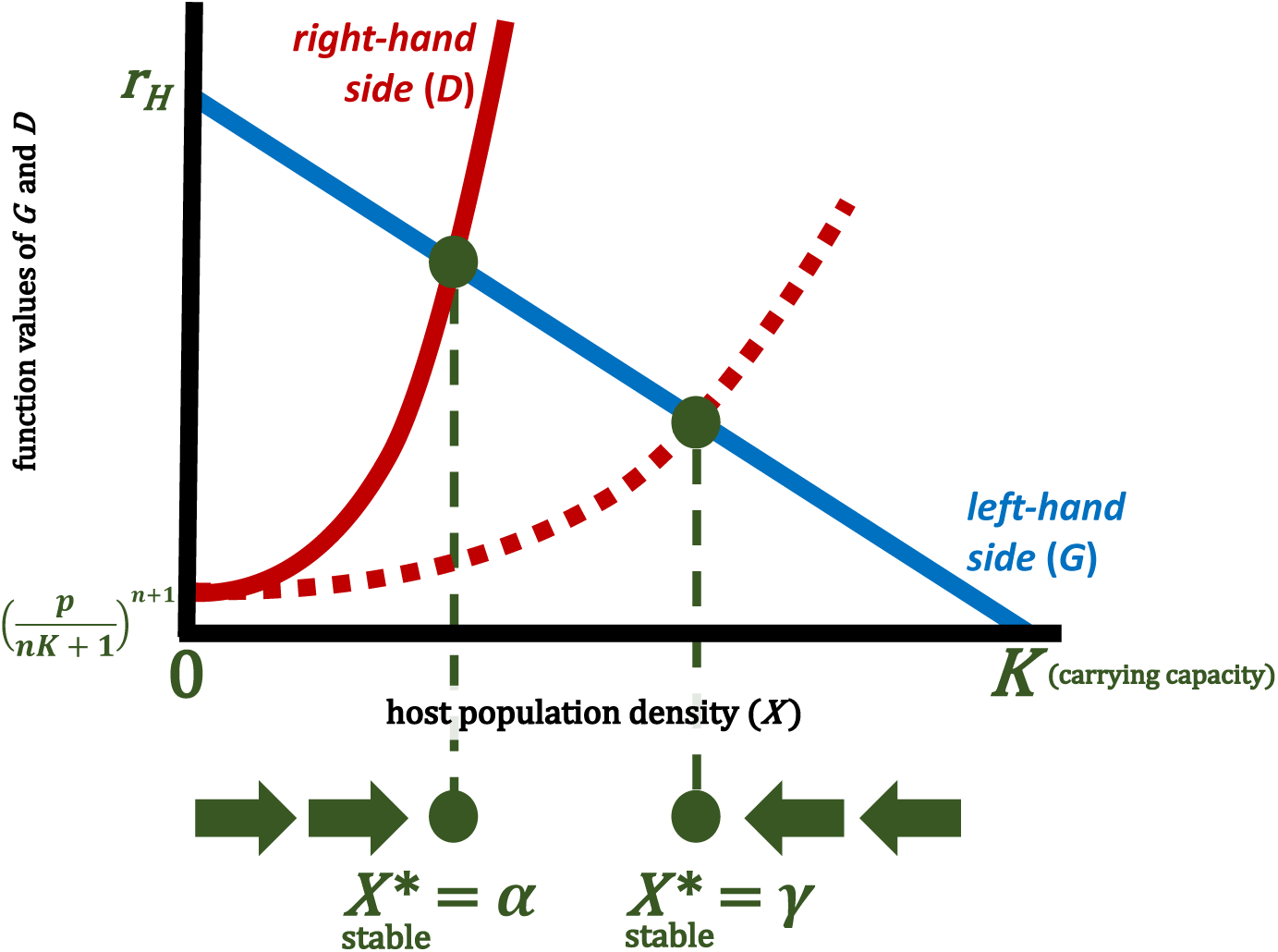
**Illustration of** **Eq.** (10), the intersection of the curves formed by the left- (blue line) and right-hand (red curve) sides of Eq. (10). Left-hand side is based on 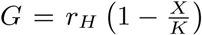, the right-hand side is based on 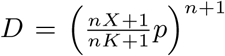. If *G* > *D* (blue line is above the red curve) then the population of host increases. If *G*<*D* (blue line is below the red curve) then the population of host decreases. The intersection of the two curves is a non-zero stable equilibrium point (*X*\*). The broken red curve represents the function 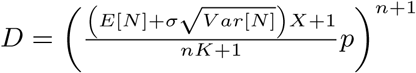.

If parasite aggregation affects the carrying capacity of the parasite population, we can replace *Y* * in Eq. 9 by an implicit equation 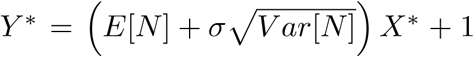.

Since 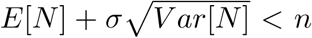, the new equilibrium state, if it exists, is *X*\* = *γ* > *α*, where *α* is the equilibrium state if *Y* * = *nX*\* +1 (Fig. 4). This relation means that aggregation limits the parasitism-driven stress affecting the host population. Sample numerical simulations of host-parasite population dynamics are shown in Fig. 5 with varying values of *σ*. The value of 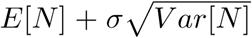 converges to *n* if the value of *σ* is increased. In the example, notice that an aggregated parasite distribution with *σ* =2 increases the parasite population rapidly but without causing too much harm to the host population (Fig. 5).

**Figure 5:**
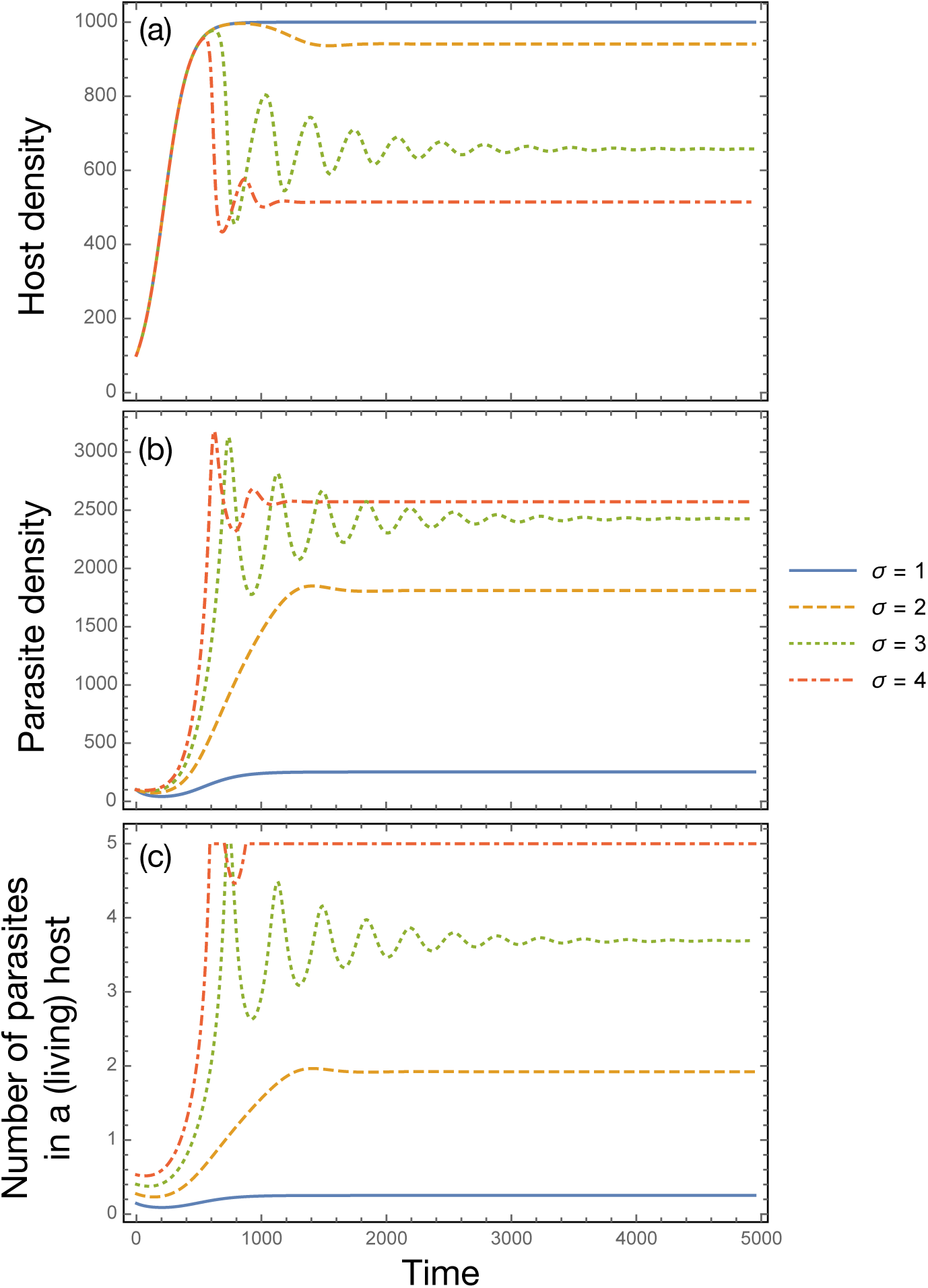
Host-parasite dynamics for different values of *σ*, the intensity of how much the variance affects the average number of parasites per host. The value of *σ* affects (A) host density and (B) parasite density. (C) The average number of parasites in a (living) host 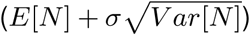 converges to the maximum possible (here *n* = 5) as the variance becomes more of a contributing factor (increase in *σ*). For low contribution of the variance (low *σ*), the hosts are typically parasite free, however increasing *σ* strengthens the host-parasite interactions leading to the typical oscillations (or an internal fixed point where both population coexist). Parameter values used in the numerics: *K* = 1000, *r*_*H*_ = 0.01, *r*_*P*_ = 0.1, and *p* = 0.8.

## Discussion

Negative-binomial distribution is commonly used in modeling macroparasite infections [46, 47]. Here, we propose a mechanistic model of parasite aggregation without initially assuming a statistical distribution. Our results show that the emergent values of the mean and variance of the macroparasite distribution indeed denote a negative-binomial. We have shown that accumulation of macroparasites (e.g., through foraging) is sufficient for aggregation to arise under a wide range of conditions (Fig. 2). The complex life cycle of parasites, a relatively large maximum tolerable parasite load *n*, and relatively moderate parasite acquisition probability *P*_*i*_ are essential factors in parasite aggregation. Thus the parasites have the opportunity to reproduce through infecting hosts but without killing many hosts. It would be rare in nature to find hosts with low tolerance *n* with parasites having high *P*_*i*_ resulting in non-aggregated parasite distribution since these hosts (as well as the parasites, if specific) would be extinct.

Our model is useful in predicting the conditions resulting in overdispersed (aggregated), underdispersed, and random parasite distributions by investigating the values of the parasite acquisition parameter *P*_*i*_. Overdispersion is observed when the variance of the parasite load in hosts is higher than the mean, while underdispersion has a mean higher than its variance. Random patterns arise if mean equals the variance. This prediction is valuable in parasitology when performing empirical studies, especially when designing statistical sampling procedures. If the parasites are aggregated in the host population, then a large number of samples is expected to be needed to select those hosts with high parasite load at the tail of the distribution [48].

The main advantage of our model compared to the classical input-output modelling framework is its simplicity without losing important biological details. Therefore, we can model host-parasite interactions using minimal models, but still, considering aggregation. The designed model framework has few variables since the effect of parasite distribution can be summarised using its moments (mean and variance). The framework supports traditional population dynamic models, such as the logistic host-parasite interaction model, which are amenable to numerical and analytic mathematical investigations. Remarkably, our assumed parameters, especially *P*_*i*_, can be directly calculated from available empirical data. Moreover, our model is more general than the stratified worm burden in terms of parasite acquisition [34]. In stratified worm burden, a host can only acquire one parasite at a time. In our model, hosts can acquire more than one parasite since compartments associated with *X*_*i*_ are defined as hosts with “at least” (not “exactly”) *i* parasite load, a scenario impossible in the stratified worm burden model.

Multiple indices are typically proposed to measure parasite aggregation. One example is the negative-binomial parameter *k*, which is equivalent to *m* in *NB*(*m, ρ*). This is defined by the equation *Var*[*N*] = *E*[*N*]+ *E*^2^[*N*]*/k*. As *k* decreases to zero (e.g., less than one but positive), the parasite distribution is said to become more aggregated. This low *k* can also infer heterogeneity in infection factors [49]. If *k* approaches infinity, the Poisson distribution results. Increase in *k* indicates a movement toward “randomness” [49, 50]. When *k* = 1, the parasite population follows a geometric distribution. Based on our results, we have identified parameters that affect the aggregation index *k*, most notably are the parameters *P*_*i*_ and *n*. This implies that *P*_*i*_ can be used as an alternative measure of aggregation that biologists can use in studying patterns of macroparasite distribution in hosts.

Given a set of parasite counts gathered from host samples, one can calculate the estimates for 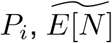 and 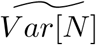. If we want to calculate the expected value of *N* ∈ {0, 1, 2, …, *n*} without assuming a large *n*, we can simply normalize the probabilities associated with *N*. That is, 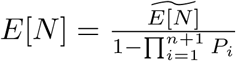. Similarly, we can do this normalization to obtain the variance of *N, Var*[*N*]. The variables and parameters *X*_*i*_, *Y*_*i*_, *P*_*i*_, *E*[*N*] and *Var*[*N*] can be functions of time, and a time series analysis to study the temporal pattern of these variables and parameters can be implemented. If *P*_*i*_’s, the host-parasite encounter probabilities, are statistically equal, we can assume a fixed *P* for each unit of time by taking the geometric mean of *P*_*i*_ (*i* ≥ 1), that is, 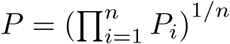.

The mortality rate due to parasitism in Eq. (3) can be modified to include the cases where a fraction of the hosts with *i*<*n* number of parasites could also die due to infection. This rate can be formulated as *bX* where *b* is as follows:

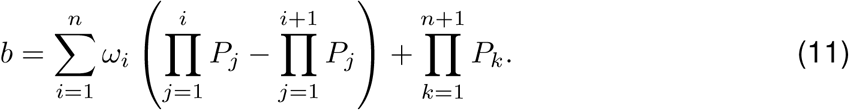

The parameter *ω*_*i*_ is the fraction of *X*_*i*_ − *X*_*i*+1_ killed by the parasites. Our conclusions from the qualitative analysis, especially given a fixed *P* = *P*_*i*_ for *i* ≥ 1, still remain true since we can suppose the mortality rate *b* as the slope of the parasite-driven host death in Fig. 3.

Here, we have determined the resulting distribution of parasites in the host population using homogeneous conditional probabilities of parasite acquisition. If the values of the acquisition probabilities become heterogeneous, such as if *P*_1_ < *P*_2_ < … < *P*_*n*_, *P*_1_ > *P*_2_ > … > *P*_*n*_ or *P*_*i*_’s are completely arbitrary, then our derived formulas to estimate *E*[*N*] and *Var*[*N*] are not applicable (e.g., Eq. (6)). However, our qualitative analysis to investigate the dynamics of Eqs. (3) and (4) could still hold, and a numerical study would then be the feasible way forward. To find appropriate formulae (if data are not available) for 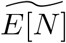 and 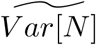 would then be a challenge for the future.

Various future studies can stem from our model design. One can include the existence of alternative, or intermediate hosts and vectors, especially that neglected tropical diseases are common due to vector-borne macroparasite infections [2]. Extending our model is possible, to include how spatial aspects and different treatment strategies affect the population dynamics of the hosts and parasites, and to infer the reasons why the value of *k* is dynamic as observed in empirical studies [51, 52, 46, 53]. Inclusion of multiple parasites [54, 55], different models of host-parasite interactions in food webs [56, 57], or the explicit inclusion of population dynamics together with host-parasite co-evolution [58, 59, 60, 61] are other possible directions. Testing if such complexities retain the observed phenomenon of parasite aggregation would then be a genuine test of the “law”.

## Acknowledgements

JFR appreciates the time spent at the Max Planck Institute for Evolutionary Biology developing the project. JFR and ELA are supported by the Institute of Mathematical Sciences and Physics, University of the Philippines Los Baños. JFR is supported by The Abdus Salam International Centre for Theoretical Physics Associate Scheme. CSG is supported by the Max Planck Society.

## Declarations

The authors declare no conflicts of interest. The funding sources had no involvement in the study design, in the analysis and interpretation of results, in the writing of the manuscript and in the decision to submit the article for publication.

## Author Contributions

JFR conceptualized the problem. JFR and CSG designed the model. JFR and ELA implemented the simulations. All authors contributed in the analysis of the model, interpretation of results, writing of the manuscript and preparation of figures.

## Data Availability

The relevant codes for generating the figures are available at https://github.com/tecoevo/parasiteaggregation

## Supplementary Material

### S.I Supplementary Information

#### SI.1.1 List of state variable and parameters

Since we have

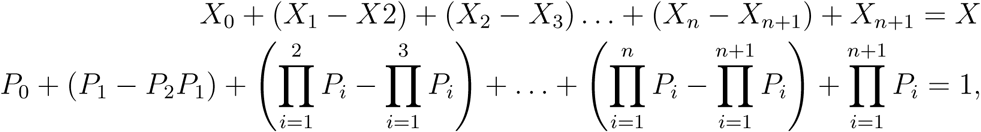

the mean of the random variable *N* +1 is

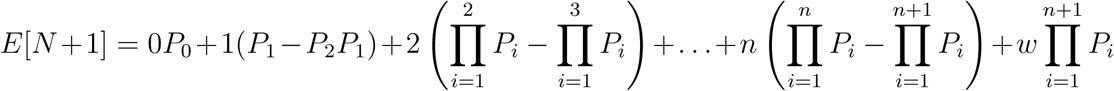

The random variable *N* +1 ∈ {0, 1, 2,…, *n, w*} (where *w* ≥ *n* + 1) represents the parasite load in living and parasitism-driven dead hosts. For simplicity, we let *w* = *n* + 1 since hosts with equal or more than *n* +1 parasites are dead due to parasitism. Let us define 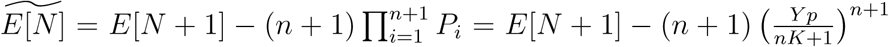, which is the approximate mean of *N* (where *N* ∈ {0, 1, 2,…, *n*}) as discussed in the main text.

**Table SI.1:**
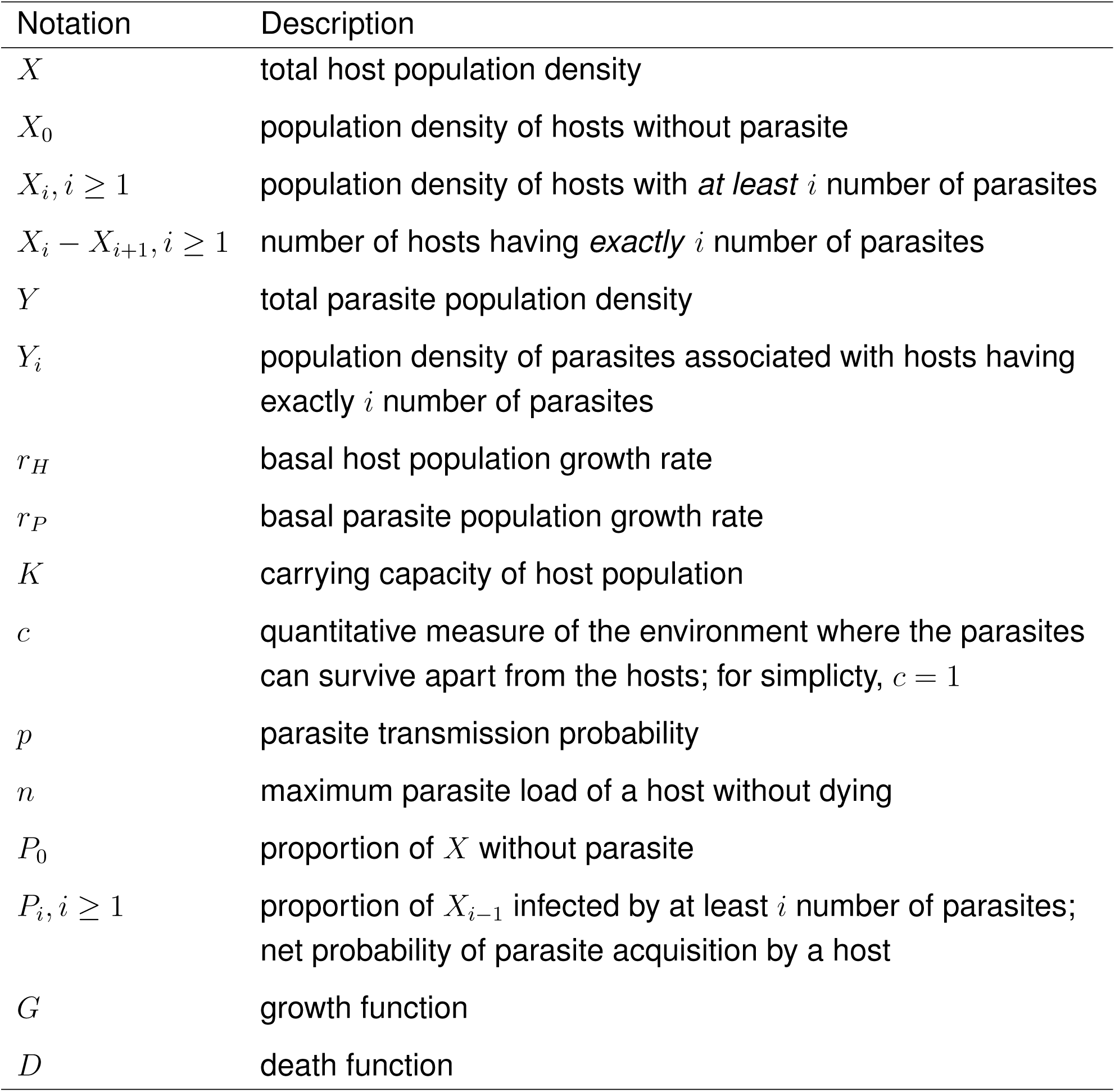
Description of the state variables and parameters. All state variables and parameters are non-negative.

**Table SI.2:**
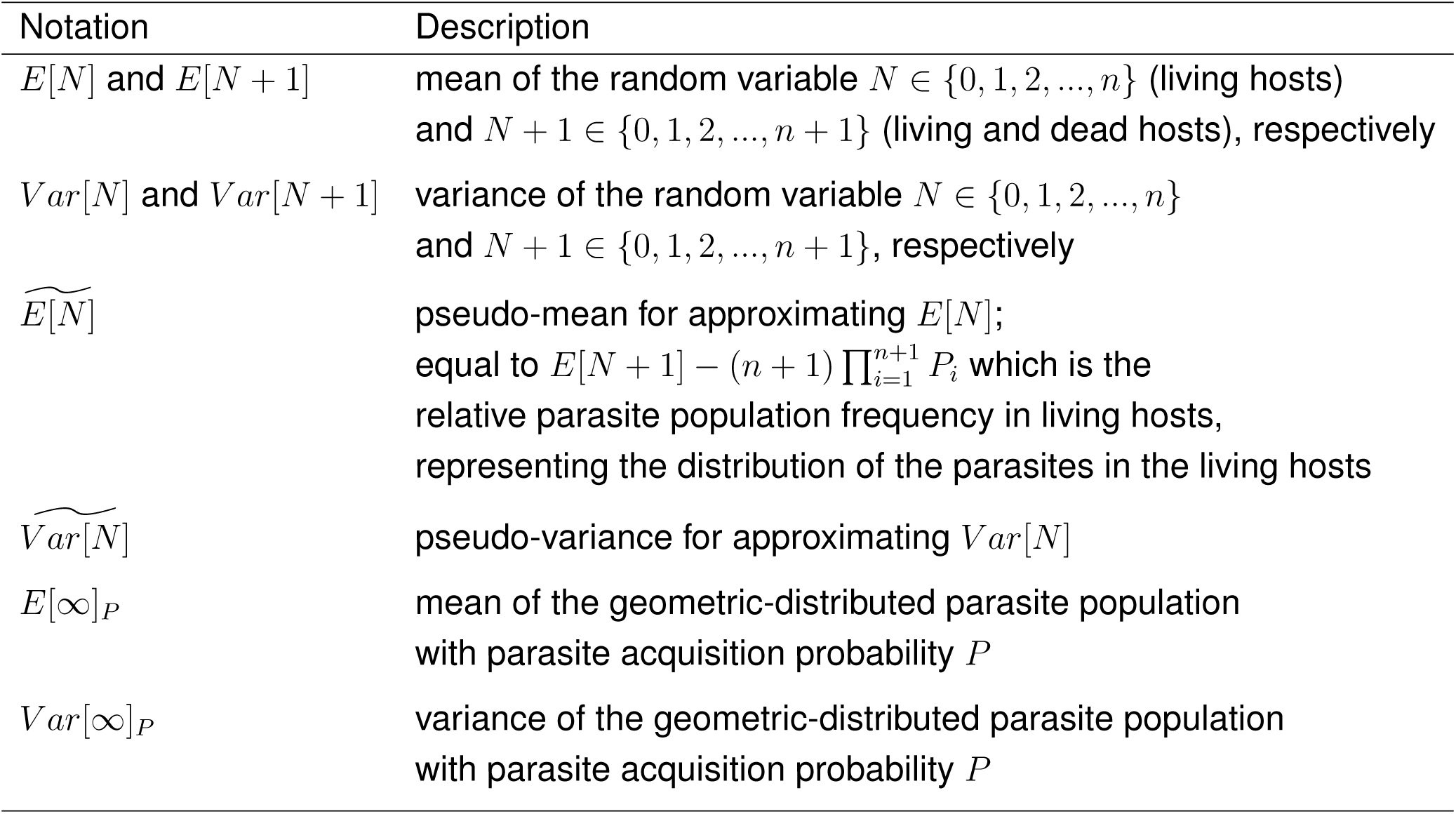
Description of the notations used in representing the mean and variance of the parasite distribution.

| Notation | Description |
| --- | --- |
| $E[N]$ and $E[N + 1]$ | mean of the random variable $N \in \{0, 1, 2, \dots, n\}$ (living hosts) and $N + 1 \in \{0, 1, 2, \dots, n + 1\}$ (living and dead hosts), respectively |
| $Var[N]$ and $Var[N + 1]$ | variance of the random variable $N \in \{0, 1, 2, \dots, n\}$ and $N + 1 \in \{0, 1, 2, \dots, n + 1\}$, respectively |
| $\widetilde{E[N]}$ | pseudo-mean for approximating $E[N]$; equal to $E[N + 1] - (n + 1) \prod_{i=1}^{n+1} P_i$ which is the relative parasite population frequency in living hosts, representing the distribution of the parasites in the living hosts |
| $\widetilde{Var[N]}$ | pseudo-variance for approximating $Var[N]$ |
| $E[\infty]_P$ | mean of the geometric-distributed parasite population with parasite acquisition probability $P$ |
| $Var[\infty]_P$ | variance of the geometric-distributed parasite population with parasite acquisition probability $P$ |

#### SI.1.2 Derivation of *A*

We assume that the parasite acquisition probability is 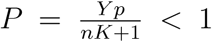. Using the geometric series, we know that,

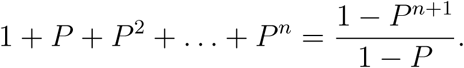

Then taking the derivative of this geometric series results in

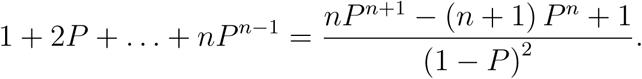

The expression for *A* is:

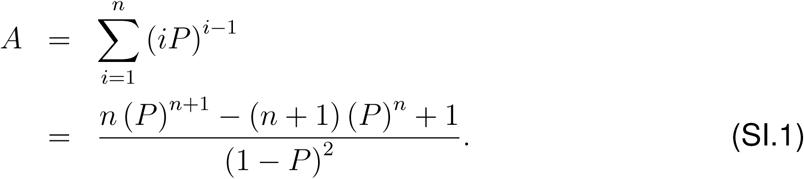

#### SI.1.3 Explicit formula for the variance

Let us define 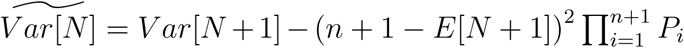 where *Var*[*N* + 1] = *E*[(*N* + 1)^2^] − *E*[*N* + 1]^2^ is the variance of the random variable *N* + 1.

With the same assumption as above for *P*, The expression for *E*[(*N* + 1)^2^] is

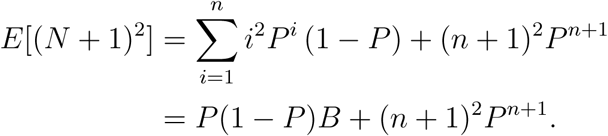

The expression for *B* can be derived using the geometric series. We know that

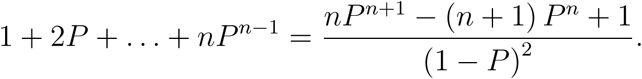

Multiplying both sides by *P*, we have

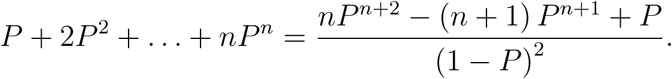

Taking the derivative of the left- and right-hand sides, we arrive at the following expression for *B*:

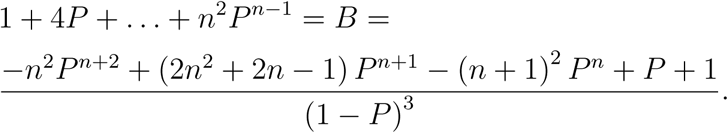

Hence, the explicit formula for 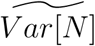 is

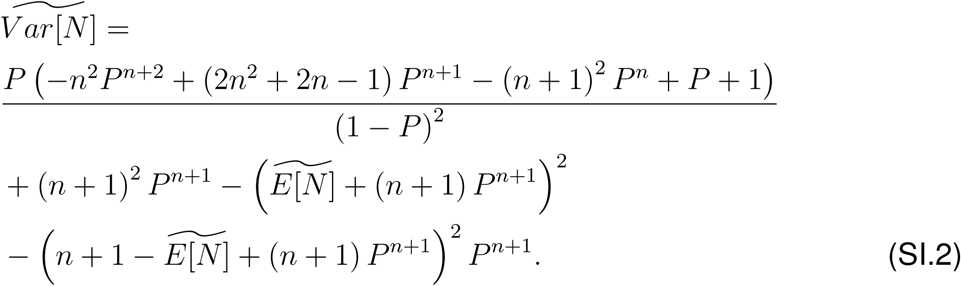

#### SI.1.4 Proof for a claim in Section “Constant host-parasite encounter probability: Population dynamics” in the main text

Here is the proof that 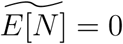 when 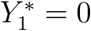 only happens if *P* = 0. Suppose *P* < 1:

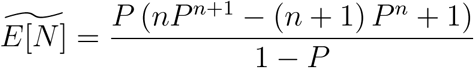

implies *P =* 0 or *nP*^*n+1*^ − (*n* + 1) *P*^*n*^ + 1 = 0. If *nP*^*n*+1^ − (*n* + 1) *P*^*n*^ + 1 = 0 then

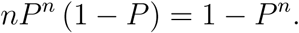

Note that 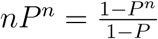 represents the geometric series:

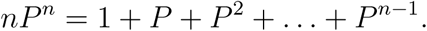

However, 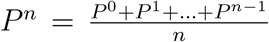 is the arithmetic average of {*P* ^*n*−1^,*P* ^*n*−2^,…,*P* ^1^,*P* ^0^}. This a contradiction since *P* ^*n*^ ∉ {*P* ^*n*−1^,*P* ^*n*−2^,…,*P* ^1^,*P* ^0^}.

Suppose *P* = 1: Note that 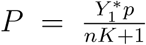 is fixed. If *P* = 1, then 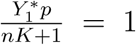 or 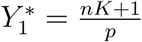. But 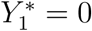 and *p* < ∞, a contradiction.

Hence, 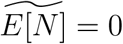 only if *P* = 0.

#### SI.1.5 Histograms given different values of *P, n* + 1= 10

**Figure SI.1:**
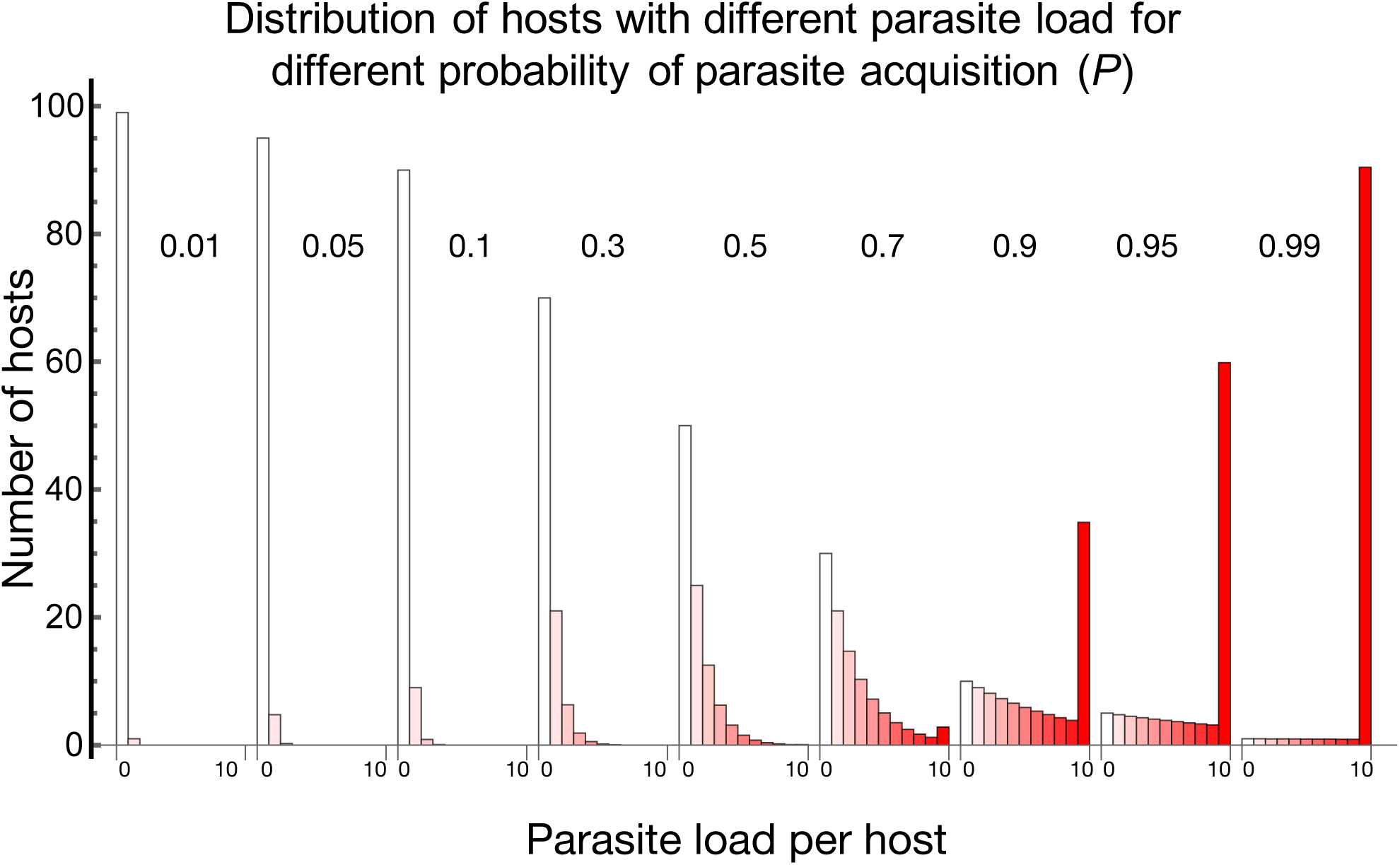
The distributions for parasite load in hosts for different parasite acquisition probabilities *P*. The distribution becomes more negatively skewed as *P* increases. Higher *P* results in high host mortality due to harbouring high parasite load. This is the reason why as *P* increases, the errors in approximating 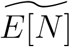 and 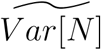 using the geometric distribution also increase (Fig. 2 in the main text).

**Figure SI.2:**
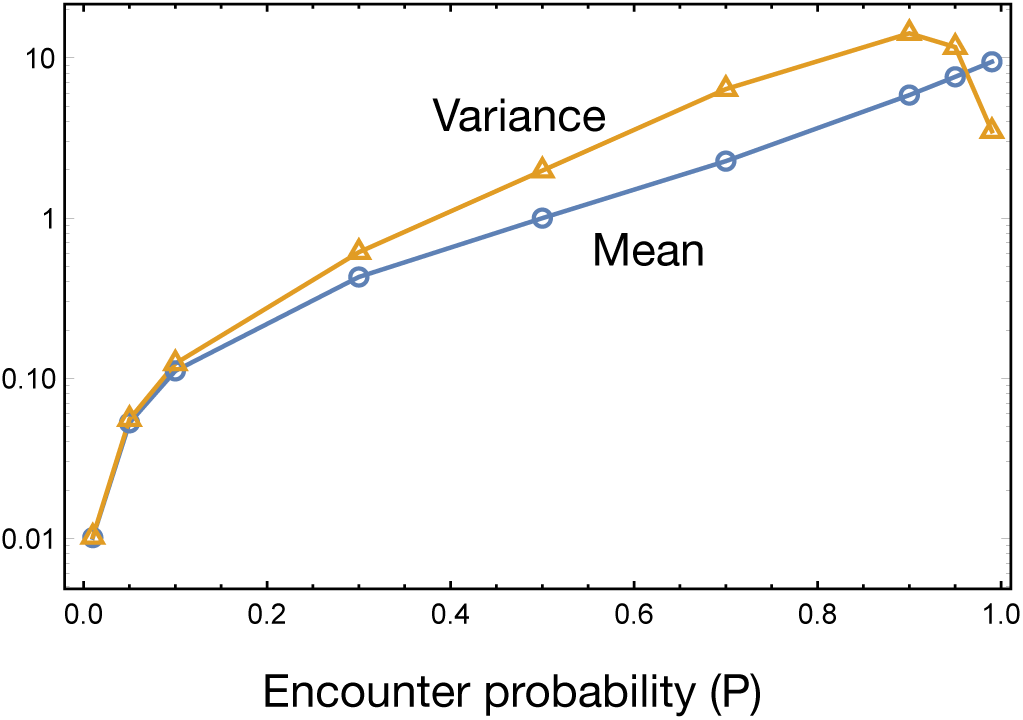
An intermediate value of *P* results in parasite aggregation (variance>mean). A value of *P* near 1 when *n* +1 = 10 results in less aggregated parasite distribution, which is possibly due to high mortality in hosts.

